# Conditional deletion of HIF-1α provides new insight regarding the murine response to gastrointestinal infection with *Salmonella* Typhimurium

**DOI:** 10.1101/2021.01.16.426940

**Authors:** Laura Robrahn, Aline Dupont, Sandra Jumpertz, Kaiyi Zhang, Christian H. Holland, Joël Guillaume, Sabrina Rappold, Vuk Cerovic, Julio Saez-Rodriguez, Mathias W. Hornef, Thorsten Cramer

**Affiliations:** Department of General, Visceral and Transplantation Surgery, RWTH University Hospital, Pauwelsstraße 30, 52074 Aachen, Germany; Institute of Medical Microbiology, RWTH University Hospital, Pauwelsstraße 30, 52074 Aachen, Germany; Institute for Computational Biomedicine, Faculty of Medicine, Heidelberg University, and Heidelberg University Hospital, Bioquant, Heidelberg, Germany; Joint Research Center for Computational Biomedicine, RWTH Aachen University Hospital, Aachen, Germany; Institute of Molecular Medicine, RWTH University Hospital, Aachen, Germany; ESCAM – European Surgery Center Aachen Maastricht, Germany and The Netherlands; Department of Surgery, Maastricht University Medical Center, Maastricht, The Netherlands; NUTRIM School of Nutrition and Translational Research in Metabolism, Maastricht University, Maastricht, The Netherlands

**Author notes:** Corresponding authors: Thorsten Cramer, Molecular Tumor Biology, General, Visceral- and Transplantation Surgery, RWTH University Hospital, Pauwelsstraße 30, 52074 Aachen, Germany, Mathias Hornef, Institute of Medical Microbiology, RWTH University Hospital, Pauwelsstraße 30, 52074 Aachen, Germany. Department of Immunology, University Medical Center Utrecht, Utrecht University, Utrecht, The Netherlands (Joël Guillaume).

## Abstract

The hypoxia-inducible transcription factor 1 (HIF-1) has been shown to ameliorate different bacterial infections through enhancement of microbial killing. While the impact of HIF-1 on inflammatory diseases of the gut has been studied intensively, its function in bacterial infections of the intestine remains largely elusive. With the help of a publicly available gene expression data set, we could infer significant activation of the HIF-1 transcription factor after oral infection of mice with *Salmonella* Typhimurium. This prompted us to apply lineage-restricted deletion of the *Hif1a* locus in mice to examine cell type-specific functions of HIF-1 in this model. We show hypoxia-independent induction of HIF-1 activity upon *Salmonella* infection in the intestinal epithelium as well as in macrophages. Surprisingly, *Hif1a* deletion in intestinal epithelial cells impacted neither disease outcome nor inflammatory activity. The conditional knockout of *Hif1a* in myeloid cells enhanced the mRNA expression of the largely pro-inflammatory chemokine *Cxcl2*, revealing a potentially inflammatory effect of HIF-1 deficiency in myeloid cells in the gut *in vivo*. Again, the disease outcome was not affected. *In vitro* HIF-1-deficient macrophages showed an overall impaired transcription of pro-inflammatory factors, however, *Salmonella* bypassed direct intracellular, bactericidal HIF-1-dependent mechanisms in a *Salmonella* pathogenicity island (SPI)-2 independent manner. Taken together, our data suggest that HIF-1 in intestinal epithelial and myeloid cells is either dispensable or compensable in the immune defense against *Salmonella* Typhimurium.

## Introduction

With drastically increasing numbers of antibiotic resistant bacteria and little novel antibiotic discoveries, new strategies to combat acute infections are urgently needed. Boosting the immune response rather than killing the pathogen directly has become a focus of investigations. The hypoxia-inducible factor (HIF)-1 functions as a cellular energy switch orchestrating energy homeostasis, metabolism, angiogenesis and evasion of apoptosis in response to low oxygen levels (hypoxia) in multiple cell types and tissues^1^. Interestingly, the heterodimeric transcription factor HIF-1 further regulates the innate immune response to invading pathogens in ways that yet remain to be understood in greater detail to evaluate its potential as a therapy target ^2,3^.

The ubiquitously and constitutively expressed HIF-1 consists of a HIF-1α and -β subunit. HIF-1α is hydroxylated by oxygen-dependent prolyl hydroxylase enzymes (PHDs) and marked for proteasomal degradation by von Hippel-Lindau tumor suppressor protein (pVHL) under normoxic or rather non-stress conditions, preventing it from otherwise binding to the HIF-1β subunit and translocating into the nucleus when oxygen levels are low ^4,5^. The closely related HIF-2α isoform has been shown to exhibit overlapping transcriptional activity but is mainly expressed in endothelial cells and bone marrow ^6,7^. Gram-positive and gram-negative bacteria as well as LPS potently stabilize HIF-1 ^2,8,9^ thereby promoting innate immune effector functions of phagocytes ^10,11^. Pharmacological stabilization of HIF-1α by inhibition of iron and oxygen-dependent PHDs utilizing iron-chelators was shown to be an effective tool against bacterial infections *in vivo* and *in vitro* ^12,13^, arguing for a potential of HIF-1 targeting for the treatment of bacterial infections.

Hypoxia, physiologically defined as oxygen concentration below 1% or partial pressure under 10 mmHg, is a hallmark of tissue inflammation, and best explained by reduced tissue perfusion combined with increased oxygen uptake by immune cells ^14^. Luminal cells of the healthy gastrointestinal epithelium are marked by physiologic hypoxia ^15,16^ which overlaps with HIF-1α stabilization. This is functionally linked to the intestinal luminal consumption of oxygen by the microbiota and the production of short-chain fatty acids which promote oxygen consumption by intestinal epithelial cells, ultimately resulting in HIF-1 protein stabilization ^17^. HIF-1 guards intestinal barrier integrity in two major ways: (i) by supporting continuous proliferation via interaction with Wnt and Notch signaling ^18,19^ and (ii) by direct barrier stabilization ^20,21^ through securing of tight junctions ^22^. While the function of HIF-1 has been extensively studied in intestinal inflammation and tumorigenesis, albeit with conflicting results ^18,20,23^, the effect of HIF-1 on bacterial infections of the intestinal tract remains largely elusive. In an elegant study, the group of Paul Beck reported a functional significance of HIF-1 for bacteria-induced intestinal injury. They showed that IEC-specific *Hif1a* knock-out mice displayed more severe intestinal injury and inflammation in response to *Clostridium difficile* infection ^24^. Furthermore, pharmacological stabilization of HIF-1α attenuated *C. difficile*-induced injury, further supporting a functional importance of the HIF-1 pathway in this setting. In line with these results, Hartmann and colleagues showed that IEC-specific *Hif1a* knock-out mice were highly susceptible to orogastric *Yersinia enterocolitica* infection. The authors linked *Y. enterocolitica*-induced HIF-1-stabilization to bacterial *s*iderophore expression, including salmochelin, a siderophore also expressed by *Salmonella* ^25^.

Through ingestion of contaminated food and water, non-typhoidal *Salmonella* represents an important cause of the world’s gastrointestinal infections ^26,27^.

*Salmonella* initially colonizes the small intestine and invades IECs, overcoming the mucosal barrier and manipulating the immune response through translocation of various effector molecules ^28–30^. It can bypass humoral immunity by hijacking phagocytes to egress from the gut into the lymphatic system or bloodstream, potentially causing life-threatening systemic infections ^31^. The above-described influence of HIF-1 on cellular immune functions as well as tight sealing of the intestinal wall therefore led us to investigate its role during gastrointestinal infections, using *Salmonella enterica* subsp. *enterica* sv. Typhimurium (*S.* Typhimurium/*S.* Tm) – a non-typhoidal strain which causes typhoid fever-like systemic infections in mice.

## Materials and Methods

### Mouse models

For constitutive deletion of *Hif1a* specifically in intestinal epithelial cells (IECs), termed *Hif1a*^IEC^, Villin-Cre transgenic mice, expressing the Cre recombinase under control of the Villin1 promotor ^32^, were crossed with mice harboring a floxed *Hif1a* locus ^33^. For inducible IEC-specific *Hif1a* deletion, VillinCre-ERT2 transgenic mice, which harbor Cre recombinase fused to the ligand binding domain of the human estrogen receptor under control of Villin1 ^34^, were crossed with *Hif1a*-floxed mice (termed *Hif1a*^IECind^). For constitutive myeloid cell-specific *Hif1a* deletion, LysMcre mice harboring Cre under control of the murine M lysozyme promotor ^35^ were crossed with *Hif1a*-floxed mice (termed *Hif1a*^MC^).

### Murine *Salmonella* infection

For *in vivo* studies, 10-week-old mice from the above-described mouse lines and their co-housed littermates were used. Mice from the inducible *Hif1a* IEC knockout line were further administered Tamoxifen (i.p., 100 mg/kg bodyweight) for five consecutive days to induce *Hif1a* deletion. For a comparable course of infection amongst experimental groups and between animals, this model requires oral streptomycin treatment of all mice before oral administration of *Salmonella* to produce a niche for bacterial colonization and invasion in the gut. Accordingly, animals were treated with streptomycin via oral gavage (20 mg/50 μL H_2_O) 48 h after the last Tamoxifen injection and/or one day before infection. The next day, animals received either 100 μL PBS (controls) or 10^8^ cfu *Salmonella* Typhimurium in 100 μL PBS, also via oral gavage ^36,37^. Four days post infection (p.i.), animals were sacrificed for organ harvest. Along control treatment or infection, mice were examined and weighed at least once daily in accordance with the approved animal protocol. Liver, spleen, mesenteric lymph nodes (MLN), small intestine (SI), cecum and colon were collected and homogenized in sterile PBS. Counts of viable *Salmonella* colony forming units (cfu) were obtained by serial dilutions and plating on LB agar plates supplemented with Ampicillin. All animal experiments (and choice of humane endpoints) were performed according to German animal protection law (TierSchG) and approved by the local animal welfare committee (Landesamt für Natur-, Umwelt- und Verbraucherschutz Nordrhein-Westfalen, Recklinghausen) under the code AZ84-02.04.2016.A491.

### Bacterial strains

For *in vivo* and *in vitro* infection experiments the Ampicillin-resistant *S.* Typhimurium strain ATCC 14028 (kindly provided by Brendan Cormack, Stanford, USA) was used. The isogenic Δ*sseB S.* Typhimurium strain ATCC 14028MvP643 and the constitutive Δ*sseBpsseB* strain ATCC 14028, Kan^R^, were kindly provided by Michael Hensel (Osnabrück, Germany) ^38^. The *E. coli* K12 strain D22 was utilized for *in vitro* use ^39^.

### IEC isolation

Intestinal epithelial cells (IECs) were isolated utilizing sections of all three parts of the small intestine. After sacrifice, intestinal sections were kept in PBS supplemented with 2% FBS (fetal bovine serum), cleaned from feces and then turned inside-out. Sections were incubated in 30 mM EDTA in PBS at 37°C. Then, IECs were detached in PBS with 2% FBS and isolated from other cells types by three sedimentation steps. The IEC pellets were snap-frozen and stored at −80°C until further preparation.

### Western Blotting

IECs (frozen pellets) or BMDMs (in petri-dishes) were lysed in nuclear extraction buffer according to Dignam and Roeder ^40^. Forty micrograms of nuclear extracts were used for SDS-PAGE and wet-transfer blotted onto a nitrocellulose membrane. Immunoblotting was performed using antibodies against HIF-1α (Cayman Chemical, Ann Arbor, USA, 1:700; 10006421), HIF-2α (Novus Biologicals, 1:1.000; NB100-122) and YY1 (Proteintech Europe, 1:1.000; 66281-1-Ig). For blocking and antibody dilution, 5% milk in TBS-T buffer was used. Densitometry was calculated using ImageJ software (NIH, Bethesda, USA) relative to YY1, and normalized to the corresponding untreated wildtype controls.

### RNA isolation and quantitative PCR

Total RNA from snap-frozen IECs was isolated with peqGOLD RNAPure™ (Erlangen, Germany) and reverse transcription was performed using the iScript cDNA Synthesis Kit (Bio-Rad, Hercules, California, USA). Following, quantitative realtime PCR (qPCR) was performed using the Applied Biosystems 7500 RealTime PCR System in a 96-well format. Reactions contained 15 ng cDNA, 0.3 μM specific primer or primer mix as indicated in manufacturers instruction at 0.1 μM and 1x Power SYBR Green Master Mix (Applied Biosystems, Bleiswijk, The Netherlands). Primer mix for *Il18* was obtained from Biomol GmbH (Hamburg, Germany). Primers (all from 5’ to 3’) against *β-actin* (F: CAC TGT CGA GTC GCG TCC, R: TCA TCC ATG GCG AAC TGG TG), *Hif1a* (F: GCT TCT GTT ATG AGG CTC ACC, R: ATG TCG CCG TCA TCT GTT AG), *Nos2* (F: AAG CCC CGC TAC TAC TCC AT, R: AAG CCA CTG ACA CTT CGC A) *Cxcl2* (F: AAG TTT GCC TTG ACC CTG AA, R: AGG CAC ATC AGG TAC GAT CC), *Cxcl5* (F: TGC CCT ACG GTG GAA GTC AT, R: AGC TTT CTT TTT GTC ACT GCC C), *Il1b* (F: CAA CCA ACA AGT GAT ATT CTC CAT G, R: GAT CCA CAC TCT CCA GCT GCA), *Il6* (F: TGA GAA AAG AGT TGT GCA ATG GC, R: GCA TCC ATC ATT TCT TTG TAT CTC TGG), *Tnfa* (F: CCA TTC CTG AGT TCT GCA AAG G, R: AGG TAG GAA GGC CTG AGA TCT TAT C), *Cramp* (F: CAG CTG TAA CGA GCC TGG TG, R: CAC CTT TGC GGA GAA GTC CA), *Lcn2* (F: ATGCACAGGTATCCTCAGGT, R: TGGCGAACTGGTTGTAGTCC) and *LysM* (F: ATG GAA TGG CTG GCT ACT ATG, R: ACC AGT ATC GGC TAT TGA TCT GA) were selected to span exon borders if possible and were validated according to the MIQE guidelines ^41^. Relative mRNA expression was calculated using the comparative delta- CT method and normalized to *β-actin* using qbase+ 3.0 (Biogazelle, Zwijnaarde, Belgium).

### Immunohistochemistry

Immunohistochemical analysis was conducted on formalin-fixed and paraffin-embedded small intestines. Sections were stained with an antibody against HIF-1α (Cayman Chemical, Ann Arbor, USA; 10006421) and counterstained with hematoxylin. Hypoxic areas were visualized using the Hypoxyprobe-1 kit (Hypoxyprobe Inc., Burlington, USA). Detailled staining procedures were previously described ^42,43^.

### Isolation and differentiation of bone marrow-derived macrophages (BMDMs)

Bone marrow from tibiae and femurs of 8- to 12-week-old WT and *Hif1a*^MC^ mice was collected and seeded onto non-TC-treated cell culture plates in Roswell Park Memorial Institute (RPMI) medium supplemented with 10% FBS overnight. Non-adherent cells were collected and cultured in RPMI (with Penicillin and Streptomycin (Pen/Strep)) supplemented with 20% FBS and 30% L929-conditioned medium and 100 U/mL penicillin, and 100 μg/mL streptomycin for 6 days for differentiation to M0 macrophages (BMDMs). Cells were then seeded overnight in RPMI (with Pen/Strep) supplemented with 10% FBS and 15% L929-conditioned medium. On assay day, cells were washed twice with PBS and medium was changed to RPMI supplemented with 2% FBS without antibiotics two hours before assay.

### Intracellular killing assay and *in vitro* infection

Bacteria were grown over night in LB medium supplemented with antibiotics as indicated. On assay day, log phase cultures were grown in LB, washed with PBS twice and then diluted in assay medium. For intracellular killing assays BMDMs were infected with a MOI (multiplicity of infection) of 10 of indicated *Salmonella* and *E. coli* strains. Plates were then briefly centrifuged to assure bacteria-cell contact. 30 minutes after infection, Gentamycin was added to cells (100 μg/mL final concentration) to kill extracellular bacteria. After one, two and four hours, cells were washed twice with PBS and then lysed with 0,0025% Triton X-100. Counts of viable bacteria/colony forming units (cfu) were obtained by serial dilutions and plating on LB agar plates. For later RNA isolation BMDMs were infected with an MOI of 1 instead and no gentamycin was added. For Lactate dehydrogenase (LDH) assay from supernatants of intracellular killing assay, were analyzed, the Pierce LDH Cytotoxicity Assay kit was used (Thermo Scientific, Waltham, Massachusetts) and Absorbance (A490 nm – A680 nm) was measured.

### Flow cytometry analyses of murine leukocyte populations in lamina propria

Peyers patches, mesenteric fat tissue and feces were removed from small intestines in HBSS supplemented with 3% FBS. Mucus was removed using nitex mesh to rinse. IECs were removed by two washes in HBSS/FBS containing 2 mM EDTA followed by collagenase VIII digestion (Sigma). Cells were then filtered using a 40 μm cell strainer and stained with the Live/Dead dye 7 Amino-actinomycin D (7AAD, Biolegend) and antibodies against CD45.2 (Biolegend, 104), MHC-II (Biolegend, M5.114.15.2), CD11c (Biolegend, N418) Ly6C (Bioegend, HK1.4), Ly6G (Biolegend, 1A8), CD103 (Biolegend, 2E7), CD11b (Biolegend, M1/70), Siglec F (BD, E50-2440) and CD64 (Biolegend, X54-5/7.1).

### Bioinformatics analysis

#### Data accessing and processing

On NCBI Gene Expression Omnibus (GEO) a publicly available transcriptomics dataset by Altmeyer *et al.* studying the effect of a *Salmonella* infection in a gastrointestinal infection (accession number: GSE19174) was identified ^44^. Raw data were downloaded with the R package GEOquery (version 2.56.0) ^45^ focusing only on the wildtype samples. Samples associated with a PARP1 knockout were discarded. Subsequently data were normalized first by removing lowly expressed genes, followed by background correction and between-array normalization using the R package vsn (version 3.56.0) ^46^. Finally, the probes were annotated with gene symbols and the expression was summarized in case of duplicated gene symbols.

#### Differential gene expression analysis

Differential gene expression analysis was performed using the R package limma (version 3.44.3) ^47^. A gene is considered significantly regulated with an absolute log fold-change (logFC) of at least 1 and false discovery rate (FDR) ≤ 0.05.

#### Transcription factor analysis

The activity of the transcription factor HIF-1 was inferred from gene expression data for each sample using the R/Bioconductor package DoRothEA (version 1.0.0, https://saezlab.github.io/dorothea/) ^48^. The activity is estimated by interrogating the expression of HIF-1 target genes. The difference in HIF-1 activity between control and infected samples was determined with a two-tailed t-test.

### Pathway analysis

We inferred the activity of Hypoxia and NF-κB for gene expression data for each sample using the R/Bioconductor packag progeny (version 1.11.1, https://saezlab.github.io/progeny/) ^49,50^. The activities are estimated by interrogating the expression of downstream affected genes instead of observing the expression of pathway members. Differences in pathway activities between control and infected samples were determined with a two-tailed t-test.

### Availability of code

The code to perform the presented bioinformatics analysis is written in R and is freely available on GitHub: https://github.com/saezlab/HIF1A-activity-salmonella.

### Statistics

Statistical analysis was performed using Prism 5.0 and 6.0 software (GraphPad Software, San Diego, California, USA). Statistical significance was determined according to one-way analysis of variance followed by Tukey post hoc test, Kruskal-Wallis test with Dunn’s post test, two-tailed t-test or Mann-Whitney U test as indicated. Differences were considered statistically significant at p < 0.05. The asterisks in the graphs indicate statistically significant changes with P values: * P < 0.05, ** P ≤ 0.01 and *** P ≤ 0.001.

## Results

### HIF-1 is highly stabilized in response to *Salmonella* in the small intestine

As infections with other enteric pathogens such as *Y. enterocolitica* result in the stabilization of HIF-1 ^2,25^, we sought to address the relevance of HIF-1 in the setting of gastrointestinal *Salmonella* infection in an oral murine model^37^. We performed a transcription factor analysis using DoRothEA ^48^ on a published gene expression data set (4 days p.i. with *Salmonella*) ^44^. This analysis convincingly demonstrated upregulation of most of HIF-1 target genes, which were summarized as an increased activity score of HIF-1 in response to *Salmonella* in the gut (Fig. 1A and B). In order to understand the upstream pathways responsible for HIF-1 transcriptional activation, we additionally applied the pathway analysis tool PROGENy to the published dataset ^49,50^. Besides hypoxia-dependent activation, this analysis suggested non-canonical HIF-1α stabilization as the NF-κB pathway was upregulated (Fig. 1C). In line with these *in silico* results, we could demonstrate robust stabilization of the epithelial HIF-1α protein accompanying mucosal tissue destruction and villus shortening at 4 days p.i. with *Salmonella* (Fig. 1D). Of note, the villus tip-pronounced hypoxia pattern vanished upon *Salmonella* infection (Fig. 1D), consistent with hypoxia-independent (non-canonical) HIF-1 stabilization in intestinal epithelial cells (IECs).

**Figure 1.**
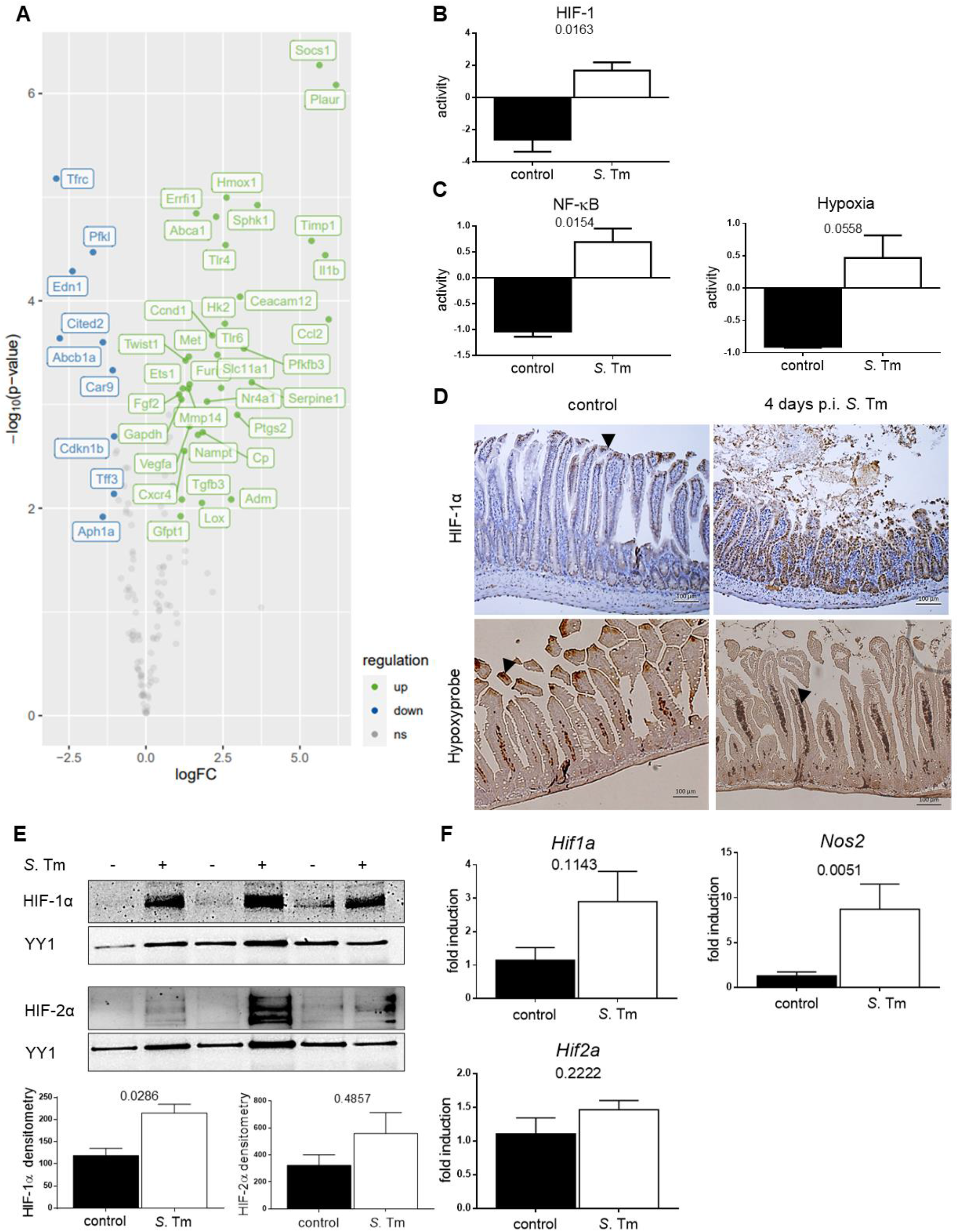
Gene expression and pathway analysis in response to hypoxia-independent stabilization of HIF-1 to *Salmonella* (*S.* Tm) in the gut. (**A**) Gene expression analysis of HIF-1 target genes after *S*. Tm infection. (**B**) Computation of HIF-1 transcriptional activity, (**C**) NF-κB and Hypoxia pathway activities upon *Salmonella* infection in comparison to uninfected controls. (**D**) Representative images of small intestines 4 days post oral PBS administration (control) or *S.* Tm infection (p.i.) stained for HIF-1α (upper panel) and Hypoxyprobe (lower panel). Arrows point towards luminal villus tips. (**E**) HIF-1α and HIF-2α western blots of nuclear extracts from IECs isolated from control or 4 days p.i. (*n*=3) with YY1 as loading control and corresponding densitometry. (**F**) qPCR analysis of *Hif1a (n=4)*, *Nos2* and *Hif2a (n=5)* mRNA expression in uninfected IECs and p.i. relative to reference gene β*-actin* and normalized to control. Data represent means with SEM. * P-values according to two-tailed t-test (B, C) or Mann-Whitney U test (F).

Small intestinal epithelial cells (IECs) were isolated for mRNA extraction and to obtain epithelial nuclear protein extracts (Fig. 1E-F). Since previous studies on chemically-induced inflammation of the gut, such as DSS colitis, hinted at an influence of HIF-2 on intestinal mucosal inflammatory processes ^51^, we examined HIF-1α as well as HIF-2α stabilization in IECs isolated from healthy and *Salmonella*-infected wild type mice. While HIF-2α protein level was enhanced in individual samples, only HIF-1α stabilization was consistently found upon infection in all infected animals (Fig. 1E). To analyze if the lack of HIF-1 accumulation could be compensated by enhanced HIF-2, we compared HIF-2α levels in HIF-1α-deficient IECs (*Hif1a*^IEC^) to WT controls and found no difference in uninfected controls or *Salmonella*-infected animals (Supplemental Fig. 1A). Next, we sought to elucidate whether the observed HIF-1 activation was transcriptional or post-translational and whether it resulted in HIF-1-dependent upregulation of HIF-1 target genes. Quantitative PCR analysis (Fig. 1F) of isolated WT IECs showed a slight upregulation of *Hif1a* and a significant upregulation of the HIF-1 target gene *Nos2* in response to *Salmonella* infection. Therefore, we conclude that epithelial HIF-1 – rather than HIF-2 – is stabilized upon *Salmonella* infection in a hypoxia-independent fashion.

### Epithelial lack of HIF-1α does not influence disease severity or systemic bacterial spread

The above results suggested a functional relevance of HIF-1 for the epithelial antimicrobial host defense against *Salmonella.* To functionally evaluate this, we used mice with a constitutive *Hif1a* knockout in IECs (termed *Hif1a*^IEC^) and assayed disease severity, inflammatory activity in the gut and bacterial spread to mesenteric lymph nodes, spleen and liver tissue. Four days after *Salmonella* infection, the indicated organs were harvested, homogenized, diluted and plated for cfu counting (Fig. 2A). Also, animals were weighed over the whole course of infection. Since slight differences in bacteria counts between individual experiments were observed and most likely caused by minor variations in the bacterial infection inoculum, bacterial numbers were normalized to those found in respective WT animals for comparison. Both wildtype litter mates and *Hif1a*^IEC^ mice exhibited significant weight loss (almost 20%), illustrating the severity of the infection. Unexpectedly, *Hif1a*^IEC^ mice showed no difference in bacterial organ counts or body weight as compared to their WT littermates. To rule out a potential functional compensation for the loss of HIF-1 in constitutively *Hif1a*^IEC^ mice, we additionally utilized a second IEC-specific knockout mouse line, where the *Hif1a* deletion in IECs was induced prior to the infection (termed *Hif1a*^IECind^). The gene deletion efficiency in both animal models – the *Hif1a*^IEC^ and *HiF1a*^IECind^ mice – was confirmed (Supplemental Figure 1B, 1C and 1D). Again, no difference in the bacterial organ load nor in weight loss was observed (Fig. 2B). Finally, we compared cytokine and chemokine and gene expression in IECs isolated from *Hif1a*^IEC^ mice in healthy controls as well as 4 days p.i. (Fig. 2C). Similarly, no significant difference was detected between both genotypes while a general upregulation was observed upon infection. Only *Cxcl5* and the HIF-1 target gene *Nos2* showed a slightly reduced upregulation in *Hif1a*-deficient IECs compared to WT.

**Figure 2.**
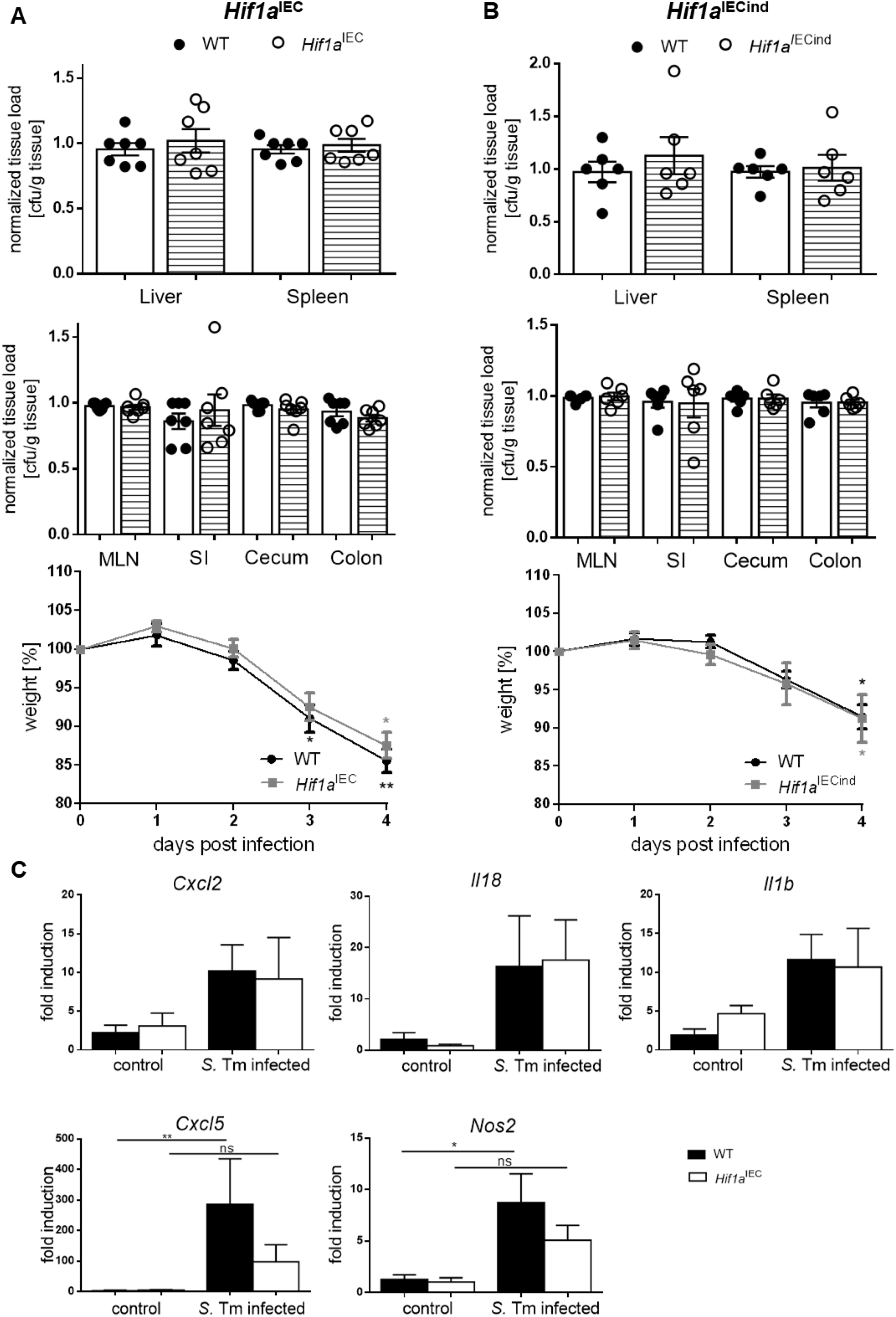
Intestinal epithelial HIF-1 loss does not affect systemic infection, weight loss or inflammatory response in a *Salmonella* infection model. Quantification of systemic *Salmonella* spread (normalized cfu (colony forming units)/g tissue) in Liver, Spleen, Mesenteric Lymph Nodes (MLN), Small Intestine (SI), Cecum and Colon and weight loss (over time course of infection) of (**A**) WT littermates and *Hif1a*^IEC^ mice harboring a constitutive HIF-1α knockout in IECs 4 days post infection *(n=7)* and (**B**) and WT littermates and Tamoxifen-inducible HIF-1α deficient (*Hif1a*^IECind^) mice (*n*=6). (**C**) mRNA expression of *Cxcl2, Il18, Il1b, Cxcl5* and *Nos2* relative to β*-actin* in WT and *Hif1a*^IEC^ IECs of control mice (*n=5*) and 4 days p.i. with *Salmonella* (*S.* Tm) (*n=7*) normalized to WT control. Data represent means with SEM. ^*^ P < 0.05; ^**^ P < 0.01; ^***^ P < 0.001 according to Kruskal-Wallis test with Dunn’s posttest (A, B, C).

### HIF-1 in myeloid cells influences the antimicrobial host response but does not influence systemic bacterial spread or disease severity

Macrophages play an essential role in the response to *Salmonella* infection by coordinating the innate and adaptive immune response through recognition of pathogen associated molecular patterns (PAMPS), leading to an antimicrobial inflammatory immune response ^52^. Of note, *Salmonella* is a facultative intracellular pathogen, and can survive within macrophages in so called *Salmonella*-containing vacuoles (SCV) bypassing the antimicrobial attack of the macrophage ^52,53^. Since the influence of HIF-1 on bactericidal functions of macrophages has been well described^10^, we explored the effect of a myeloid cell-specific *Hif1a* loss on systemic infection and disease outcome as well as mucosal inflammation. Unexpectedly, evaluation of organ load and body weight (Fig. 3A) did not show significant differences between wildtype and *Hif1a*^MC^ mice. mRNA expression analysis of IECs isolated from infected wildtype and *Hif1a*^MC^ animals showed significantly higher expression of the chemokine *Cxcl2* while *Cxcl5* expression was slightly reduced in infected *Hif1a*^MC^ mice (Fig. 3B). To further characterize the infiltration of immune cells in the mucosal tissue of *Hif1a*^IEC^ and *Hif1a*^MC^ mice, FACS analysis of isolated lamina propria cells was performed. However, no difference in the composition of various leukocyte subsets in both mouse lines compared to their wild type littermates was detected (Supplemental Fig. 2).

**Figure 3.**
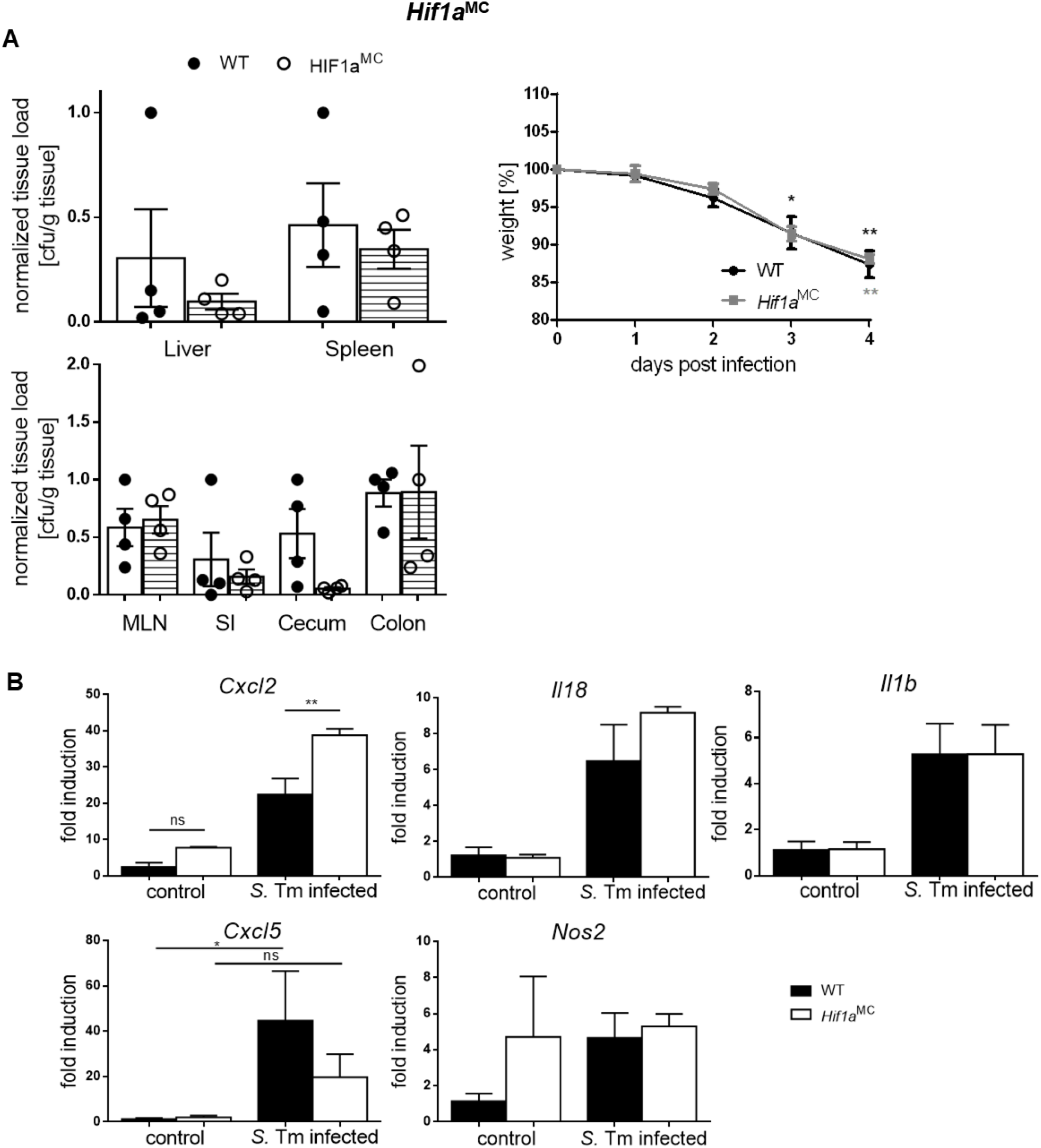
Myeloid cell specific HIF-1α deletion affects *Cxcl2* expression in IECs but not systemic *Salmonella* infection. (**A**) Quantification of systemic *Salmonella* spread/organ cfu (colony forming units) after 4 days of infection and quantification of weight loss during time course of infection of WT littermates and *Hif1a*^MC^ mice harboring a HIF-1α knockout in myeloid cells (*n=4*). (**B**) mRNA expression of *Cxcl2, Il18, Il1b, Cxcl5* and *Nos2* relative to β*-actin* in IECs from WT and *Hif1a*^MC^ uninfected controls (*n=3*) and 4 days p.i. with *Salmonella* (*S.* Tm) (*n=4*) normalized to WT control. Data represent means with SEM. ^*^ P < 0.05; ^**^ P < 0.01; ^***^ P < 0.001 according to Kruskal-Wallis test with Dunn’s posttest.

### HIF-1 is highly stabilized in macrophages upon *Salmonella* infection and influences the inflammatory response

Given the well-established supportive role of HIF-1 for microbial killing of phagocytes we were surprised that neither *Hif1a*^MC^ nor *Hif1a*^IEC^/Hif1a^IECind^ mice exhibited an altered bacterial spread after *Salmonella* infection. As macrophages represent the first line of defense against *Salmonella* ^31^, we next sought to characterize the functional importance of HIF-1α for macrophages *in vitro*. Bone marrow-derived macrophages (BMDMs) were generated from wildtype mice and infected with *Salmonella* at a multiplicity of infection (MOI) of 10 (Fig. 4A). Western blot analysis showed robust stabilization of HIF-1α upon *Salmonella* infection. To further elucidate HIF-1’s impact on the transcriptional response to *Salmonella*, gene expression analysis in BMDMs of *Hif1a*^MC^ mice and their wildtype littermates were performed. The expression of genes involved in direct bactericidal or inflammatory functions of phagocytes and in the response to *Salmonella* was compared (Fig. 4B and C). Of note, while the overall mRNA expression of pro-inflammatory mediators upon infection in HIF-1α deficient macrophages was lower, this functional inactivation of had no significant influence on the *Salmonella*-induced regulation of the pro-inflammatory mediators *Il1b, Il6* and *Tnfa* (Fig. 4B), *Lcn2* (inhibiting intracellular *Salmonella* growth due to iron chelation ^54^), *Nos2*, various chemokines and the antimicrobial peptide cathelin-related antimicrobial peptide (CRAMP) ^55^ (Fig. 4C).

**Figure 4.**
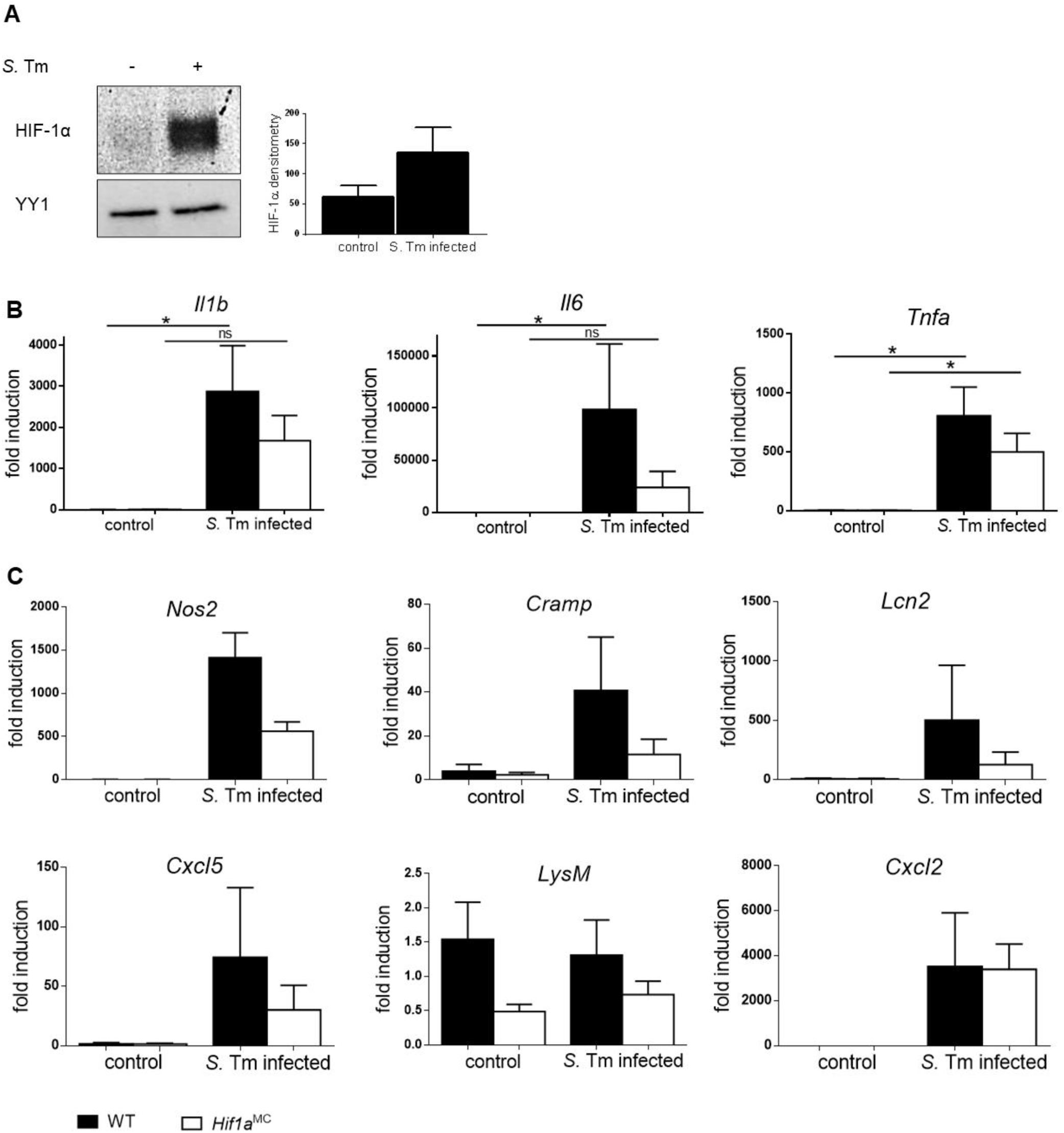
*Salmonella* (*S.* Tm)-dependent HIF-1 stabilization alters the transcription of Cytokines, AMPs and Chemokines in Macrophages. (**A**) Representative image of HIF-1α western Blot and densitometry relative to loading control YY1 (*n=3*) of control and *Salmonella* infected bone marrow derived macrophages (BMDMs), MOI 10. Relative mRNA expression of WT and *Hif1a*^MC^ BMDMs in uninfected controls and 4 hours p.i. with *Salmonella* (MOI 1) of (**B**) Cytokines *Il1b, Il6* and *Tnfa*, (**C**) AMPs (anti-microbial peptides) and Chemokines *Nos2*, *Cramp, Cxcl2, Cxcl5, Lcn2* and *LysM* (Lysozyme) relative to β*-actin* (*n*=5, *Nos2: n*=3) and normalized to WT control. Data represent means with SEM. ^*^ P < 0.05; ^**^ P < 0.01; ^***^ P < 0.001 according to Mann-Whitney U test (A) or Kruskal-Wallis test with Dunn’s posttest (B, C).

### HIF-1 does affect *E. coli* but not*Salmonella* killing by macrophages

In order to reconcile our observation with the known role of HIF-1 for the antimicrobial activity in macrophages ^11,13^, we performed comparative intracellular killing assays using *Salmonella* and the non-pathogenic commensal bacterium *E. coli*. Both WT and *Hif1a*^MC^ BMDMs exhibited equally and significantly reduced intracellular *Salmonella* counts over the time course of 4 hours of infection (Fig. 5A). In sharp contrast, while WT macrophages killed a significant percentage of intracellular *E. coli*, *Hif1a*^MC^ cells failed to do so (Fig. 5B). To investigate whether macrophages were killing intracellular bacteria or were lysed by them in turn, we measured LDH (lactate dehydrogenase) activity in the cell culture supernatant as an indicator for cell death (Fig. 5C). LDH levels in both WT and *Hif1a*-deficient BMDMs were elevated upon *Salmonella* infection already after 2 hours, while LDH levels upon *E. coli* infection were still as low as in the control. This suggested that *Salmonella* induced cell death of a notable number of infected BMDMs in this setting. Intracellularly, macrophages can kill bacteria through an oxidative burst utilizing ROS (reactive oxygen species) and RNS (reactive nitrogen species). Unlike *E. coli*, *Salmonella* is able to evade and bypass this antimicrobial host defense mechanism ^56^. *Salmonella* converts the phagosome through expression of a type III (T3SS) secretion system and translocation of effector proteins encoded by the *Salmonella* pathogenicity island 2 (SPI-2) into so-called *Salmonella*-containing vacuoles (SCV). Here, Salmonella survives and even replicates. We used a *sseB* mutant strain (Δ*sseB*) lacking a functional SPI-2 T3SS and preventing SCV formation (Supplemental Figure 3) as well as the complemented Δ*sseBpsseB* strain ^38^. However, no significant influence of the presence of HIF-1α on the number of intracellular SPI-2 T3SS deficient *Salmonella* was noted. These results do not support a significant influence of the SPI-2 T3SS on the resistance to HIF-1 induced bacterial killing in macrophages.

**Figure 5.**
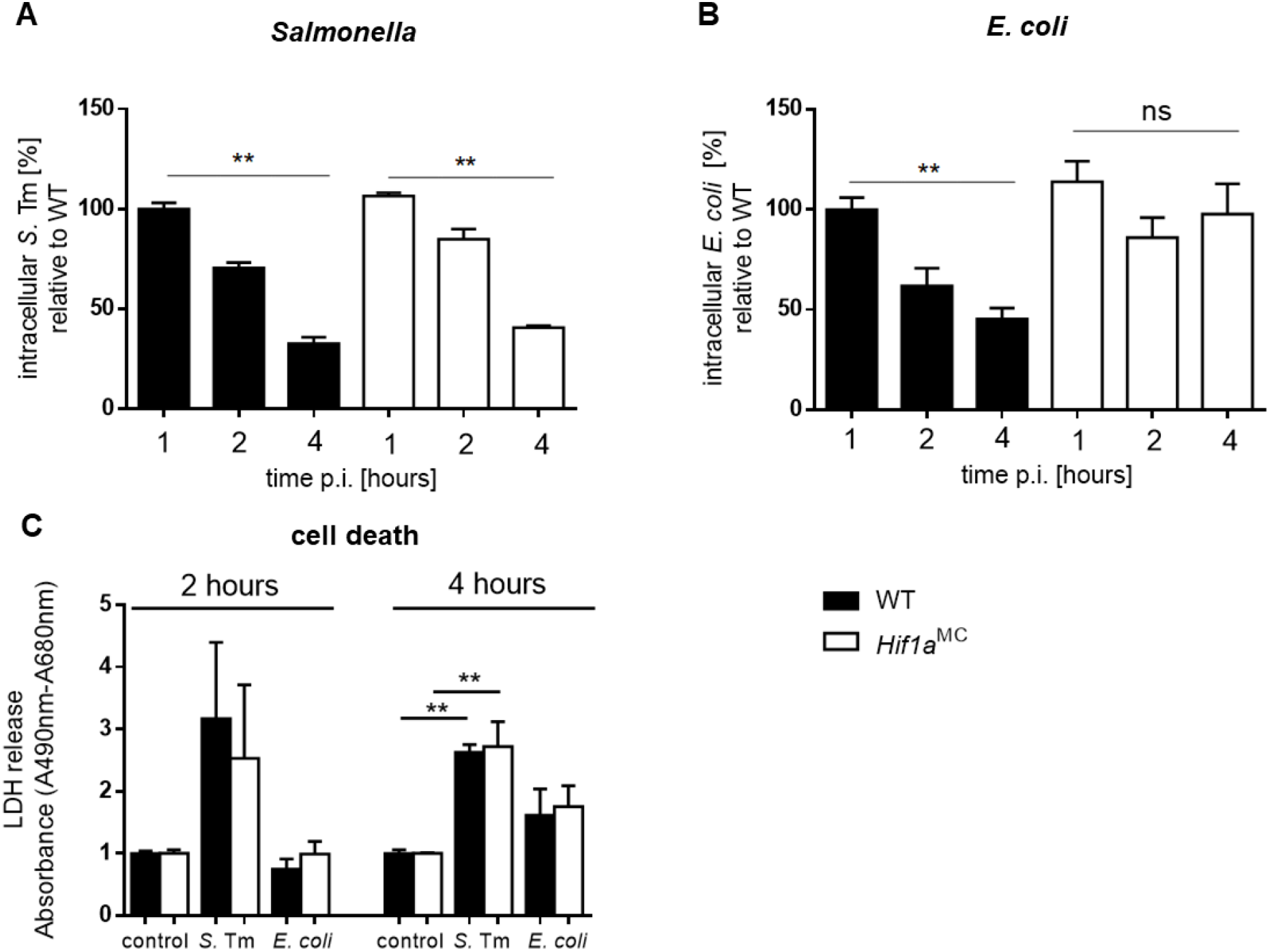
*E. coli* not *Salmonella* intracellular survival in macrophages is HIF-1 dependent. **A**) Intracellular survival of *Salmonella* in bone marrow derived WT littermates and HIF-1α deficient (*Hif1a*^MC^) macrophages (BMDM) over time course of 4 hours (MOI 10; *n*=3). Extracellular bacteria were killed by addition of medium containing Gentamycin after 30 min of incubation. (**B**) Intracellular survival of *E. coli* in WT and *Hif1a*^MC^ BMDMs over time course of infection (*n*=4). (**C**) Cell death analysis measured by lactate dehydrogenase (LDH) release into surrounding medium of controls, *Salmonella* and *E. coli* infected BMDMs after 2 and 4 hours (Absorbance A490nm-A680nm). Data represent means with SEM. ^*^ P < 0.05; ^**^ P < 0.01; ^***^ P < 0.001 according to Kruskal-Wallis test with Dunn’s posttest (A, B, C).

## Discussion

Originally identified as a master regulator of the cellular response to hypoxia, HIF-1 emerged as a central regulator of immune cell functions ^57^. Utilizing the same mechanisms as in the hypoxic setting, it shifts cellular metabolism and changes the transcription of a broad spectrum of genes and therefore influences immune functions of epithelial and endothelial cell types ^13,58^ as well as phagocytes ^10,11^ making it a potential tool to fight various bacterial and non-bacterial infections. The activation of macrophages with different inflammatory stimuli such as succinate ^59^ or bacteria (as shown for *Mycobacterium tuberculosis*) leads to a shift from the TCA cycle to glycolysis, HIF-1 stabilization and further macrophage activation ^60–62^. HIF-1 stabilization facilitates the induction of pro-inflammatory cytokines such as Il-1β and of additional mechanisms controlling intracellular bacteria in particular ^59,63^. In this study we addressed the role of HIF-1 on intestinal infections with *Salmonella,* utilizing genetically modified mice harboring either a myeloid-(*Hif1a*^MC^) or an intestinal epithelial cell-specific (*Hif1a*^IEC^) *Hif1a* knockout ^32,33,35^.

Although HIF-1α was robustly upregulated in IECs as well as macrophages in response to *Salmonella* infection, its deletion did not change the overall disease outcome in the respective infection models. No difference between wildtype and knockout mice was observed comparing weight loss as well as systemic organ spread of *Salmonella* in both settings. Since mice with a deletion of *Hif1a* in IECs develop normally, we further analyzed whether these results could be due to a compensatory mechanism and utilized a tamoxifen-inducible IEC-specific *Hif1a* knockout mouse strain (*Hif1a*^IECind^) ^33,34^. However, systemic bacteria spread and weight loss were comparable to the constitutive IEC-specific *Hif1a* knockout line.

Different research groups have reported HIF-1 to impact enteric inflammatory diseases – especially IBD and DSS-induced colitis – and thereby to play a critical role in intestinal tumorigenesis ^18,64^. Surprisingly, the *Hif1a* deletion in IECs did not alter the mRNA expression of genes encoding different cytokines, chemokines and AMPs potentially involved in the inflammatory response to bacterial pathogens. Only a slight decrease of *Cxcl5* and *Nos2* was observed upon *Hif1a* deletion. On the contrary, *Hif1a*^MC^ mice showed increase in *Cxcl2* mRNA expression upon *Salmonella* infection – hinting at a potential anti-inflammatory phenotype of HIF-1 possessing myeloid cells in this setting. Also in *vitro, Hif1a*-deficient macrophages showed globally lower mRNA expression levels of multiple effectors involved in the specific response to *Salmonella*. Yet again, intracellular *Salmonella* survival was comparable between *Hif1a*^MC^ and wildtype cells. Two macrophage mechanisms with central importance for killing of intracellular pathogens are the production of ROS and RNS. The TCA cycle intermediates succinate and fumarate have been shown to inhibit PHD function and induce ROS formation leading to robust HIF-1 stabilization ^59,65^. NO species also inhibit PHDs and stabilize HIF-1 additionally through S-nitrosylation ^66^. However, a *Salmonella* (Δ*SseB*) mutant which is more susceptible to ROS due to its inability to form *Salmonella* containing vacuoles (SCV) intracellularly ^38^ again showed comparable intracellular survival in both wildtype and *Hif1a* deficient macrophages, while – consistent with recent studies ^13^ – *E. coli* survival was positively impacted by *Hif1a* deficiency. Therefore, we conclude that *Salmonella* bypasses the ROS-response in a different, possibly yet unknown, way.

Several studies have shown inducible NO synthase (NOS2) and therefore NO (nitric oxide) to be of functional relevance for HIF-1-dependent immunity and the induction of a positive feedback-loop which drives the inflammatory macrophage response ^61^. However, chronologically the induction of NOS2 and reactive nitrogen species follows the induction of ROS at later stages of a *Salmonella* infection ^52^. In all examined mouse lines, a weight loss of almost 20% was observed, demonstrating a severe course of the infection, but potentially obscuring a HIF-1-dependent phenotype in the very final stages of a *Salmonella*-infection model portraying survival.

*Salmonella* has further been shown to evade the RNS response by limiting NOS2 substrates ^67^. *Salmonella* also exhibit ROS/RNS detoxification enzymes such as SodC (Superoxide dismutase) ^68^ on the one hand as well as PhoPQ ^69^, a factor crucial for evasion of nitrosative stress and intracellular survival, on the other hand. They could therefore avoid or alter the main HIF-1 immune response of phagocytes in general. *Salmonella* such as other intracellular bacteria like *Chlamydophila pneumoniae* ^70^ and *Francisella tularensis* ^71^ – which actively impair HIF-1 stabilization or possess HIF-1-degrading virulence factors – might therefore not be susceptible to HIF-1 dependent immunity and rather reprogram the ROS and RNS response. Taken together, our findings indicate that the functional importance of HIF-1 for bacterial killing depends on the pathogen under investigation. For a diverse range of bacteria, from *E. coli* ^13^ to MRSA ^12^, HIF could represent a therapeutic target but potentially not for all. It is important to note that our study did not address the effects of (pharmacological) stabilization of HIF-α. It surely is possible that this approach can result in protection against Salmonella, as has been shown for other infectious agents ^12,13^. Evidently, a more detailed understanding of HIF-1 immunity and bacterial adaption mechanisms of professional intracellular pathogens like *Salmonella* towards it are necessary to fully evaluate HIF-1’s potential as a therapy target.

## Abbreviations

HIF: Hypoxia-inducible factor
*S*. Tm: *Salmonella* Typhimurium
PHD: Prolyl Hydroxylase
IEC: Intestinal epithelial cells
NOS2: NO Synthetase-2
CXCL2: C-X-C motif chemokine 2
SPI-2: Salmonella pathogenicity island 2
BMDM: Bone marrow-derived macrophages
IBD: Inflammatory bowel disease
RNS: Reactive nitrogen species

## Authorship

M.H.W. and T.C. jointly supervised this study. L.R., M.W.H. and T.C. conceptualized the study. L.R., A.D., S.J., K.Z., J.G., S.R. and V.C. performed experiments and analyzed data. C.H. and J.S.R. predicted pathway activities based on microarray data. L.R. and T.C. wrote the original manuscript draft. L.R., A.D., S.J., K.Z., C.H., J.G., V.C., J.S.R., M.W.H. and T.C. reviewed and edited the manuscript.

## Acknowledgements

Research in the Cramer lab was supported by grants from the Deutsche Forschungsgemeinschaft (CR 133/2-1 until 2-4 and CR 133/3-1). Laura Robrahn is supported by a START grant from the Medical Faculty of the RWTH Aachen University. The excellent technical assistance of Johanna Roth is highly appreciated. This work has partly been presented in poster form at “Novel Concepts in Innate Immunity 2019” in Tübingen, Germany.

---

**Supplemental Figure 1.**
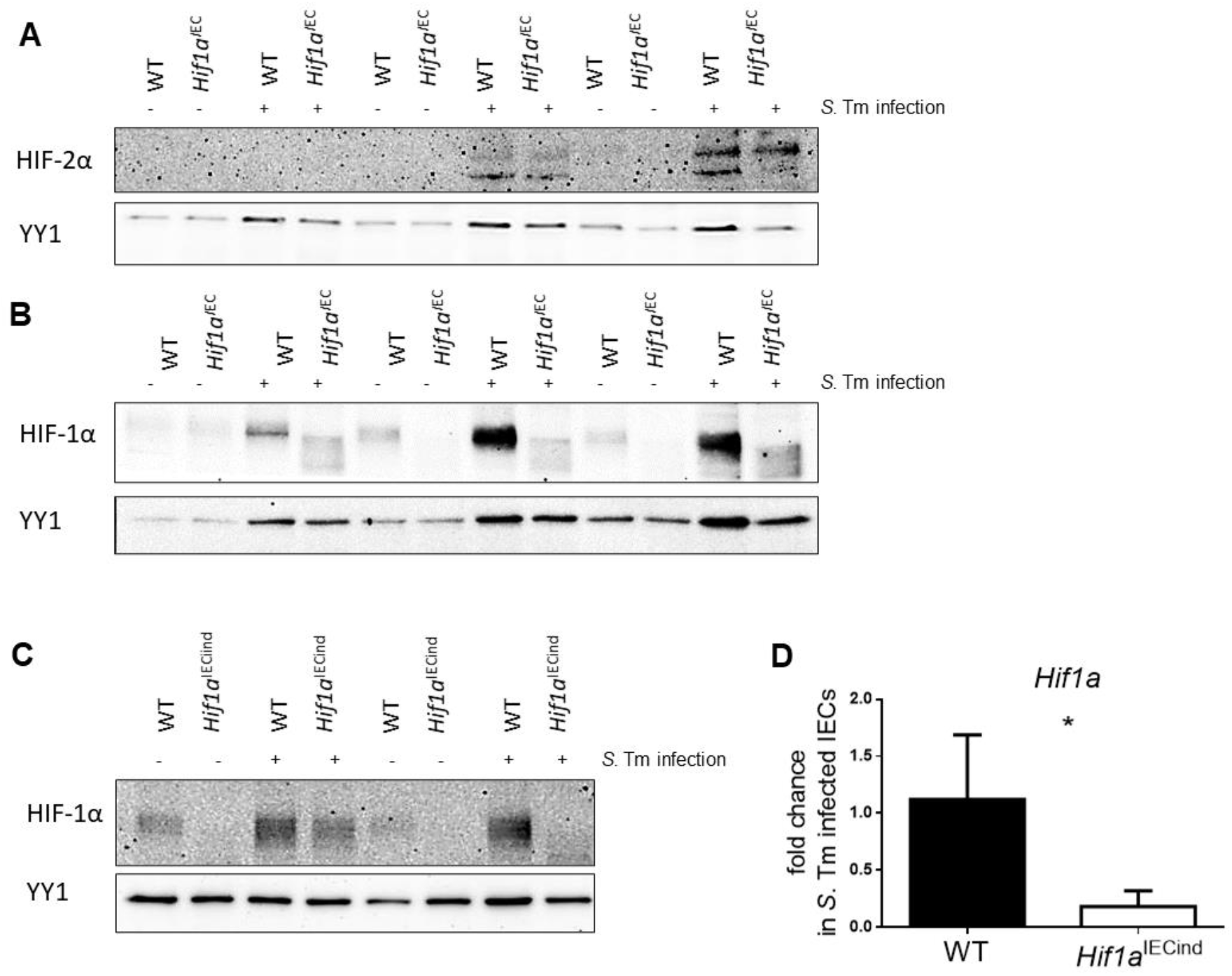
HIF-1 knockout efficiency in constitutive and Tamoxifen-inducible VillinCre mice. (**A**) HIF-2α and (**B**) HIF-1α western blots of nuclear extracts of IECs from of uninfected controls and infected (4 days p.i.) WT littermates and *Hif1a*^IEC^ mice (*n*=3) and (**C**) WT and *Hif1a*^IECind^ animals harboring a Tamoxifen-inducible Hif-1α knockout in IECs (n=2). (**D**) *Hif1a* mRNA expression relative to β*-actin* upon *Salmonella* (*S*. Tm) infection in WT and *Hif1a*^IECind^ IECs (*n*=3). Data represent means with SEM. ^*^ P < 0.05; ^**^ P < 0.01; ^***^ P < 0.001 according to Mann-Whitney U test (D).

**Supplemental Figure 2.**
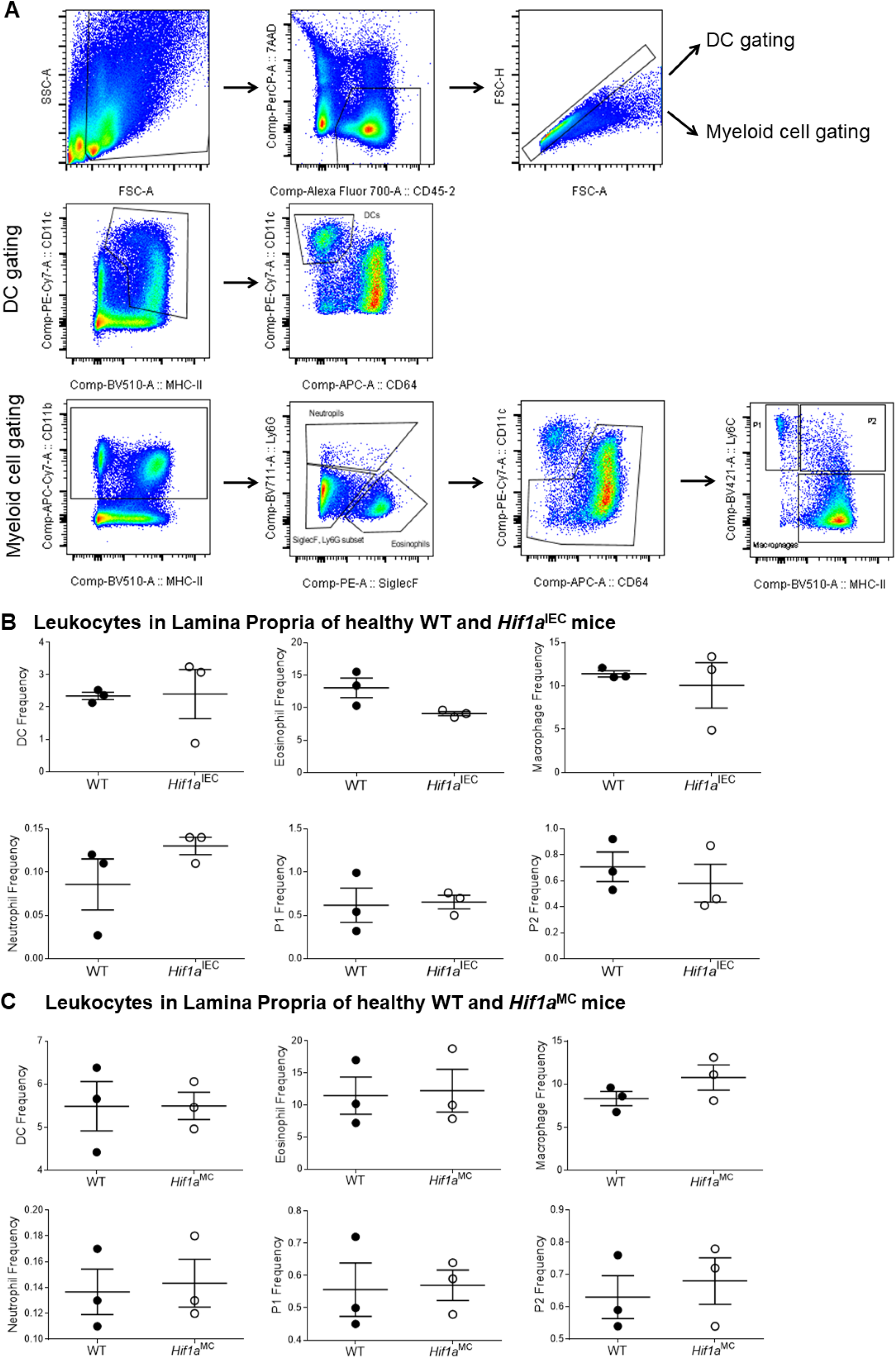
FACS-based Leukocyte Quantification in Lamina propria of uninfected WT littermates, *Hif1a* ^IEC^ and *Hif1a*^MC^ animals. (**A**) Gating strategy for DC and myeloid cell gating. Counts of Neutrophils, including P1 and P2 Neutrophils, Eosinophils and Macrophages in the small intestinal Lamina propria of (**B**) *Hif1a*^IEC^ and (**C**) *Hif1a*^MC^ mice and their WT littermates (*n*=3). Data represent means with SEM. ^*^ P < 0.05; ^**^ P < 0.01; ^***^ P < 0.001 according to Mann-Whitney U test (B, C).

**Supplemental Figure 3.**
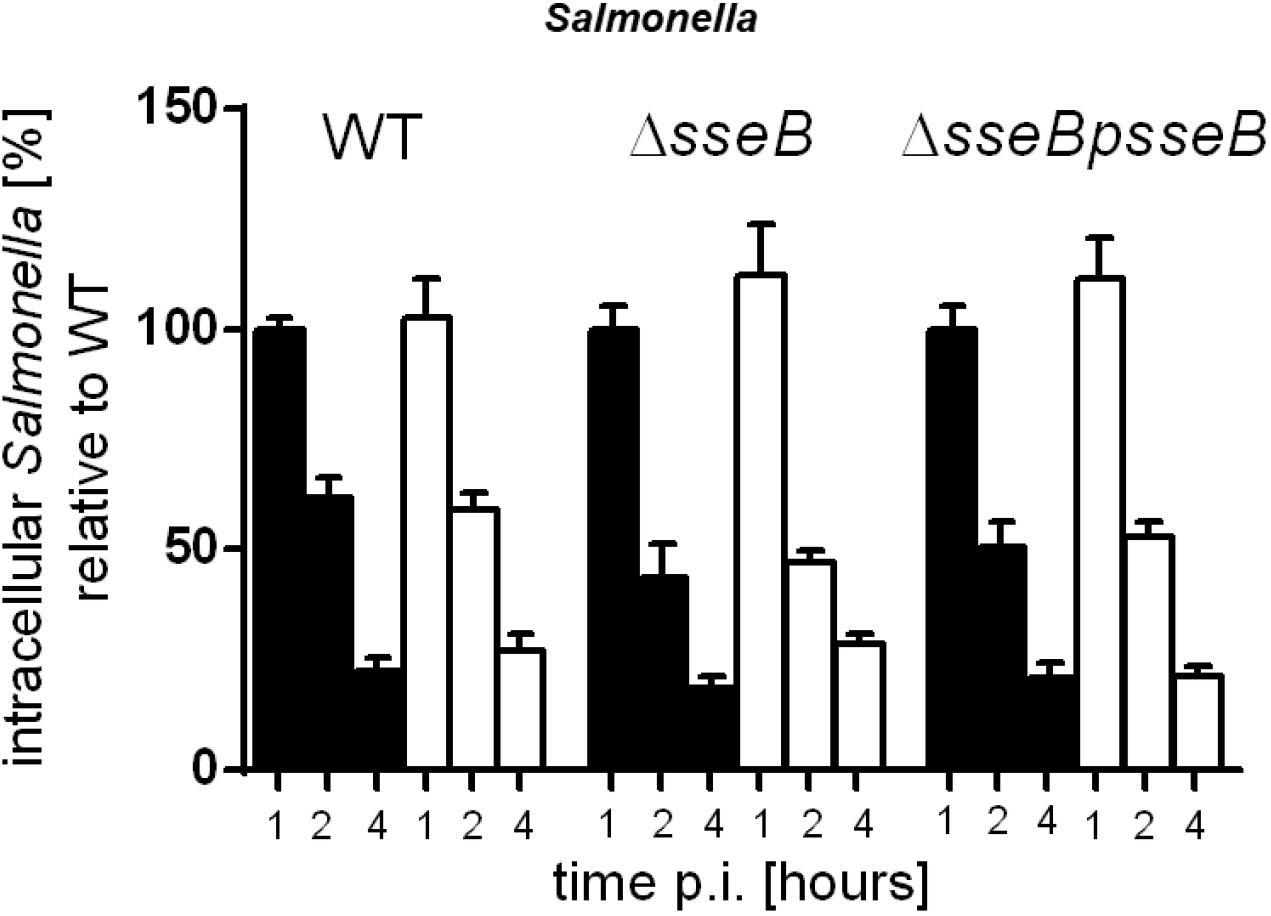
SPI-2 deficiency in *Salmonella* does not interfere with intracellular HIF-1-dependent bactericidal functions of Macrophages. (**A**) Intracellular survival of *Salmonella* in bone marrow derived WT and HIF-1α deficient (*Hif1a*^MC^) macrophages within 4 hours of infection utilizing wildtype *Salmonella*, a Δ*sseB* mutant and the complemented strain Δ*sseBpsseB* (*n=3*). Data represent means with SEM. ^*^ P < 0.05; ^**^ P < 0.01; ^***^ P < 0.001 according to one-way analysis of variance followed by Tukey post hoc test (A, B).

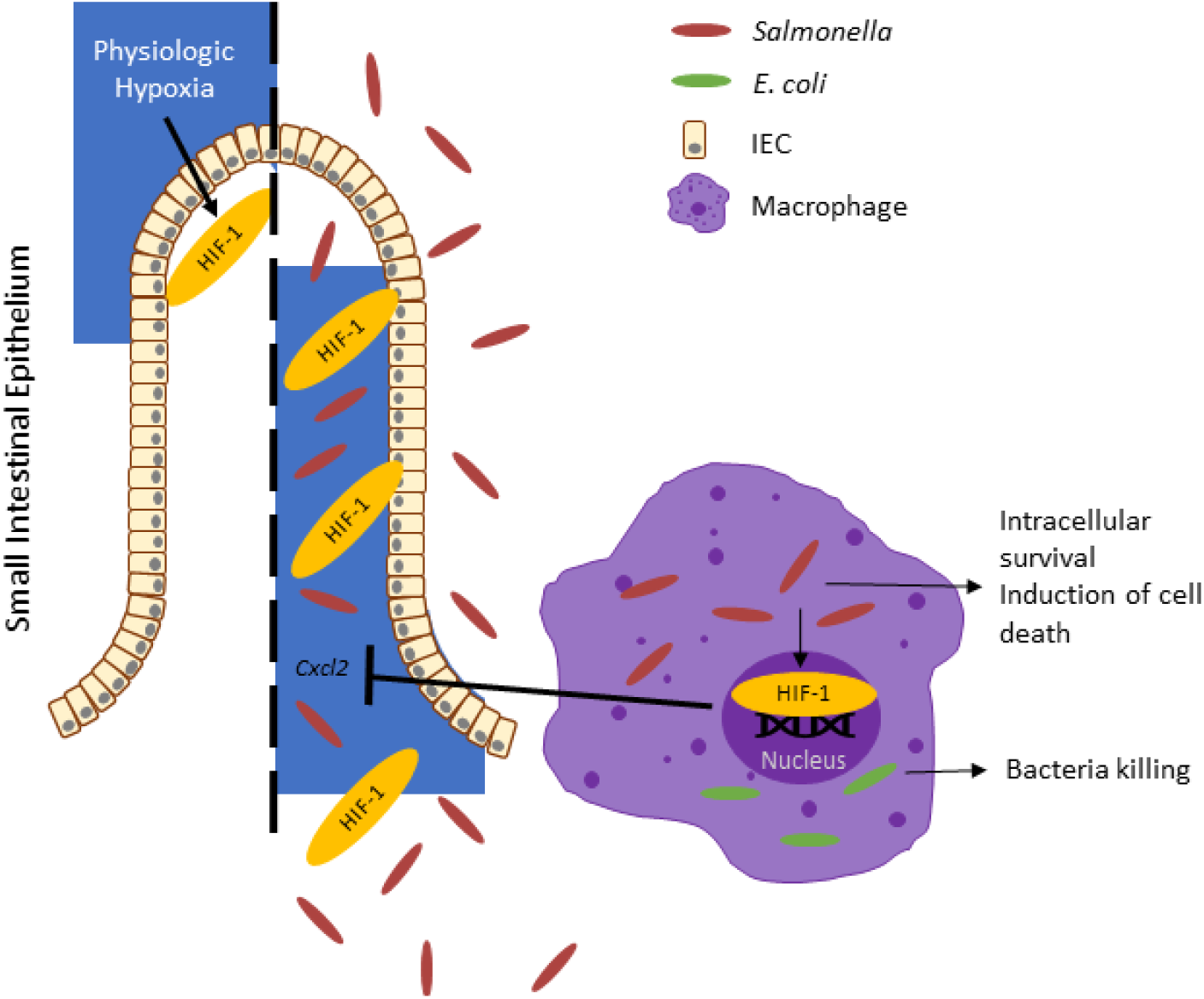
Graphical Abstract. *Salmonella* potently activates HIF-1 in the intestinal epithelium and in macrophages in a hypoxia-independent manner but bypasses its immune response.

## References

1. Choudhry H, Harris AL. Advances in Hypoxia-Inducible Factor Biology. Cell Metabolism. 2018;27:281–298.

2. Werth N, Beerlage C, Rosenberger C, et al. Activation of hypoxia inducible factor 1 is a general phenomenon in infections with human pathogens. PLoS One. 2010;5:1–12.

3. Nizet V, Johnson RS. Interdependence of hypoxic and innate immune responses. Nature Reviews Immunology. 2009;9:609–617.

4. Semenza GL, Wang GL. A nuclear factor induced by hypoxia via de novo protein synthesis binds to the human erythropoietin gene enhancer at a site required for transcriptional activation. Mol Cell Biol. 1992;12:5447–54.

5. Yu F, White SB, Zhao Q, et al. HIF-1α binding to VHL is regulated by stimulus-sensitive proline hydroxylation. Proc Natl Acad Sci U S A. 2001;98:9630–9635.

6. Ema M, Taya S, Yokotani N, et al. A novel bHLH-PAS factor with close sequence similarity to hypoxia-inducible factor 1α regulates the VEGF expression and is potentially involved in lung and vascular development. Proc Natl Acad Sci U S A. 1997;94:4273–4278.

7. Talks KL, Turley H, Gatter KC, et al. The expression and distribution of the hypoxia-inducible factors HIF-1α and HIF-2α in normal human tissues, cancers, and tumor-associated macrophages. Am J Pathol. 2000;157:411–421.

8. Rius J, Guma M, Schachtrup C, et al. NF-κB links innate immunity to the hypoxic response through transcriptional regulation of HIF-1α. Nature. 2008;453:807–811.

9. Brooks AC, Menzies-Gow N, Bailey SR, et al. Endotoxin-induced HIF-1α stabilisation in equine endothelial cells: Synergistic action with hypoxia. Inflamm Res. 2010;59:689–698.

10. Cramer T, Yamanishi Y, Clausen BE, et al. HIF-1alpha is essential for myeloid cell-mediated inflammation. Cell. 2003;112:645–57.

11. Peyssonnaux Datta, V., Cramer, T., Doedens, A., Theodarakis, E. A., Gallo, R. L., Hurtado-Ziola, N., Nizet, V., and Johnson, R. S. C. HIF-1 alpha expression regulates the bacterial capacity of phagocytes. J Clin Invest. 2005;115:1806–1815.

12. Okumura CYM, Hollands A, Tran DN, et al. A new pharmacological agent (AKB-4924) stabilizes hypoxia inducible factor-1 (HIF-1) and increases skin innate defenses against bacterial infection. J Mol Med. 2012;90:1079–1089.

13. Lin AE, Beasley FC, Olson J, et al. Role of Hypoxia Inducible Factor-1α (HIF-1α) in Innate Defense against Uropathogenic Escherichia coli Infection. PLOS Pathog. 2015;11:e1004818.

14. Silver IA. Tissue PO2 changes in acute inflammation. Adv Exp Med Biol. 1977;94:769–774.

15. He G, Shankar RA, Chzhan M, et al. Noninvasive measurement of anatomic structure and intraluminal oxygenation in the gastrointestinal tract of living mice with spatial and spectral EPR imaging. Proc Natl Acad Sci U S A. 1999;96:4586–4591.

16. Albenberg L, Esipova T V., Judge CP, et al. Correlation between intraluminal oxygen gradient and radial partitioning of intestinal microbiota. Gastroenterology. 2014;147:1055–1063.e8.

17. Kelly CJ, Zheng L, Campbell EL, et al. Crosstalk between microbiota-derived short-chain fatty acids and intestinal epithelial HIF augments tissue barrier function. Cell Host Microbe. 2015;17:662–671.

18. Rohwer N, Jumpertz S, Erdem M, et al. Non-canonical HIF-1 stabilization contributes to intestinal tumorigenesis. Oncogene. 2019;38:5670–5685.

19. Gustafsson M V., Zheng X, Pereira T, et al. Hypoxia requires Notch signaling to maintain the undifferentiated cell state. Dev Cell. 2005;9:617–628.

20. Robinson A, Keely S, Karhausen J, et al. Mucosal Protection by Hypoxia-Inducible Factor Prolyl Hydroxylase Inhibition. Gastroenterology. 2008;134:145–155.

21. Tambuwala MM, Cummins EP, Lenihan CR, et al. Loss of Prolyl Hydroxylase-1 Protects Against Colitis Through Reduced Epithelial Cell Apoptosis and Increased Barrier Function. Gastroenterology. 2010;139:2093–2101.

22. Saeedi BJ, Kao DJ, Kitzenberg DA, et al. HIF-dependent regulation of claudin-1 is central to intestinal epithelial tight junction integrity. Mol Biol Cell. 2015;26:2252–2262.

23. Cummins EP, Seeballuck F, Keely SJ, et al. The hydroxylase inhibitor dimethyloxalylglycine is protective in a murine model of colitis. Gastroenterology. 2008;134:156–65.

24. Hirota SA, Fines K, Ng J, et al. Hypoxia-Inducible Factor Signaling Provides Protection in Clostridium difficile-Induced Intestinal Injury. Gastroenterology;139. Epub ahead of print 2010. DOI: 10.1053/j.gastro.2010.03.045.

25. Hartmann H, Eltzschig HK, Wurz H, et al. Hypoxia-Independent Activation of HIF-1 by Enterobacteriaceae and Their Siderophores. Gastroenterology. 2008;134:756–767.e6.

26. Coburn B, Grassl GA, Finlay BB. Salmonella, the host and disease: A brief review. Immunology and Cell Biology. 2007;85:112–118.

27. Majowicz SE, Musto J, Scallan E, et al. The Global Burden of Nontyphoidal Salmonella Gastroenteritis. Clin Infect Dis. 2010;50:882–889.

28. Larock DL, Chaudhary A, Miller SI. Salmonellae interactions with host processes. Nature Reviews Microbiology. 2015;13:191–205.

29. Que F, Wu S, Huang R. Salmonella pathogenicity Island 1(SPI-1) at work. Curr Microbiol. 2013;66:582–587.

30. Zhang K, Riba A, Nietschke M, et al. Minimal SPI1-T3SS effector requirement for Salmonella enterocyte invasion and intracellular proliferation in vivo. PLOS Pathog. 2018;14:e1006925.

31. Sun Y, Reid B, Ferreira F, et al. Infection-generated electric field in gut epithelium drives bidirectional migration of macrophages. PLoS Biol. 2019;17:e3000044.

32. Madison BB, Dunbar L, Qiao XT, et al. cis elements of the villin gene control expression in restricted domains of the vertical (crypt) and horizontal (duodenum, cecum) axes of the intestine. J Biol Chem. 2002;277:33275–33283.

33. Ryan HE, Poloni M, Mcnulty W, et al. Hypoxia-inducible Factor-1 Is a Positive Factor in Solid Tumor Growth. 2000.

34. el Marjou F, Janssen K-P, Chang BH-J, et al. Tissue-specific and inducible Cre-mediated recombination in the gut epithelium. Genesis. 2004;39:186–93.

35. Clausen BE, Burkhardt C, Reith W, et al. Conditional gene targeting in macrophages and granulocytes using LysMcre mice. Transgenic Res. 1999;8:265–77.

36. Meynell GG, Subbaiah T V. Antibacterial mechanisms of the mouse gut. I. Kinetics of infection by Salmonella typhi-murium in normal and streptomycin-treated mice studied with abortive transductants. Br J Exp Pathol. 1963;44:197–208.

37. Barthel M, Hapfelmeier S, Quintanilla-Martínez L, et al. Pretreatment of Mice with Streptomycin Provides a Salmonella enterica Serovar Typhimurium Colitis Model That Allows Analysis of Both Pathogen and Host. Infect Immun. 2003;71:2839–2858.

38. Beuzon CR, Banks G, Deiwick J, et al. pH-dependent secretion of SseB, a product of the SPI-2 type III secretion system of Salmonella typhimurium. Mol Microbiol. 1999;33:806–816.

39. Normark S, Boman HG, Matsson E. Mutant of Escherichia coli with anomalous cell division and ability to decrease episomally and chromosomally mediated resistance to ampicillin and several other antibiotics. J Bacteriol. 1969;97:1334–1342.

40. Carey MF, Peterson CL, Smale ST. Dignam and Roeder nuclear extract preparation. Cold Spring Harb Protoc. 2009;4:pdb.prot5330.

41. Bustin SA, Benes V, Garson JA, et al. The MIQE Guidelines: Minimum Information for Publication of Quantitative Real-Time PCR Experiments. Epub ahead of print 2009. DOI: 10.1373/clinchem.2008.112797.

42. Rohwer N, Lobitz S, Daskalow K, et al. HIF-1α determines the metastatic potential of gastric cancer cells. Br J Cancer. 2009;100:772–781.

43. Daskalow K, Rohwer N, Raskopf E, et al. Role of hypoxia-inducible transcription factor 1α for progression and chemosensitivity of murine hepatocellular carcinoma. J Mol Med. 2010;88:817–827.

44. Altmeyer M, Barthel M, Eberhard M, et al. Absence of poly(ADP-Ribose) polymerase 1 delays the onset of Salmonella enterica serovar typhimurium-induced gut inflammation. Infect Immun. 2010;78:3420–3431.

45. Sean D, Meltzer PS. GEOquery: A bridge between the Gene Expression Omnibus (GEO) and BioConductor. Bioinformatics. 2007;23:1846–1847.

46. Huber W, Von Heydebreck A, Sültmann H, et al. Variance stabilization applied to microarray data calibration and to the quantification of differential expression. In: Bioinformatics. Oxford University Press:96–104.

47. Ritchie ME, Phipson B, Wu D, et al. Limma powers differential expression analyses for RNA-sequencing and microarray studies. Nucleic Acids Res. 2015;43:e47.

48. Garcia-Alonso L, Holland CH, Ibrahim MM, et al. Benchmark and integration of resources for the estimation of human transcription factor activities. Genome Res. 2019;29:1363–1375.

49. Schubert M, Klinger B, Klünemann M, et al. Perturbation-response genes reveal signaling footprints in cancer gene expression. Nat Commun. 2018;9:1–11.

50. Holland CH, Szalai B, Saez-Rodriguez J. Transfer of regulatory knowledge from human to mouse for functional genomics analysis. Biochim Biophys Acta - Gene Regul Mech. 2020;1863:194431.

51. Ramakrishnan SK, Shah YM. Role of Intestinal HIF-2α in Health and Disease. Annu Rev Physiol. 2016;78:301–325.

52. Gogoi M, Shreenivas MM, Chakravortty a-c D, et al. Hoodwinking the Big-Eater to Prosper: The Salmonella-Macrophage Paradigm Journal of Innate Immunity. Epub ahead of print 2018. DOI: 10.1159/000490953.

53. Alpuche-Aranda CM, Racoosin EL, Swanson JA, et al. Salmonella stimulate macrophage macropinocytosis and persist within spacious phagosomes. J Exp Med. 1994;179:601–608.

54. Schaible UE, Kaufmann SHE. Iron and microbial infection. Nature Reviews Microbiology. 2004;2:946–953.

55. Rosenberger CM, Gallo RL, Finlay BB. Interplay between antibacterial effectors: A macrophage antimicrobial peptide impairs intracellular Salmonella replication. Proc Natl Acad Sci U S A. 2004;101:2422–2427.

56. Figueira R, Holden DW. Functions of the Salmonella pathogenicity island 2 (SPI-2) type III secretion system effectors. Microbiology. 2012;158:1147–1161.

57. Palazon A, Goldrath AW, Nizet V, et al. Review HIF Transcription Factors, Inflammation, and Immunity. Immunity. 2014;41:518–528.

58. Leire E, Olson J, Isaacs H, et al. Role of hypoxia inducible factor-1 in keratinocyte inflammatory response and neutrophil recruitment. J Inflamm (United Kingdom);10. Epub ahead of print 2013. DOI: 10.1186/1476-9255-10-28.

59. Tannahill GM, Curtis AM, Adamik J, et al. Succinate is an inflammatory signal that induces IL-1β through HIF-1α. Nature. 2013;496:238–242.

60. Rodríguez-Prados J-C, Través PG, Cuenca J, et al. Substrate Fate in Activated Macrophages: A Comparison between Innate, Classic, and Alternative Activation. J Immunol. 2010;185:605–614.

61. Braverman J, Sogi KM, Benjamin D, et al. HIF-1α Is an Essential Mediator of IFN-γ– Dependent Immunity to Mycobacterium tuberculosis. J Immunol. 2016;197:1287–1297.

62. Gleeson LE, Sheedy FJ, Palsson-McDermott EM, et al. Cutting Edge: Mycobacterium tuberculosis Induces Aerobic Glycolysis in Human Alveolar Macrophages That Is Required for Control of Intracellular Bacillary Replication. J Immunol. 2016;196:2444–2449.

63. Knight M, Stanley S. HIF-1α as a central mediator of cellular resistance to intracellular pathogens. Curr Opin Immunol. 2019;60:111–116.

64. Robrahn L, Jiao L, Cramer T. Barrier integrity and chronic inflammation mediated by HIF-1 impact on intestinal tumorigenesis. Cancer Lett;490. Epub ahead of print 2020. DOI: 10.1016/j.canlet.2020.07.002.

65. Sudarshan S, Sourbier C, Kong H-S, et al. Fumarate Hydratase Deficiency in Renal Cancer Induces Glycolytic Addiction and Hypoxia-Inducible Transcription Factor 1α Stabilization by Glucose-Dependent Generation of Reactive Oxygen Species. Mol Cell Biol. 2009;29:4080–4090.

66. Li F, Sonveaux P, Rabbani ZN, et al. Regulation of HIF-1α Stability through S-Nitrosylation. Mol Cell. 2007;26:63–74.

67. Lahiri A, Das P, Chakravortty D. Arginase modulates Salmonella induced nitric oxide production in RAW264.7 macrophages and is required for Salmonella pathogenesis in mice model of infection. Microbes Infect. 2008;10:1166–1174.

68. Uzzau S, Bossi L, Figueroa-Bossi N. Differential accumulation of Salmonella[Cu, Zn] superoxide dismutases SodCI and SodCII in intracellular bacteria: correlation with their relative contribution to pathogenicity. Mol Microbiol. 2002;46:147–156.

69. Bourret TJ, Liu L, Shaw JA, et al. Magnesium homeostasis protects Salmonella against nitrooxidative stress. Sci Rep;7. Epub ahead of print December 1, 2017. DOI: 10.1038/s41598-017-15445-y.

70. Rupp J, Gieffers J, Klinger M, et al. *Chlamydia pneumoniae* directly interferes with HIF-1α stabilization in human host cells. Cell Microbiol. 2007;9:2181–2191.

71. Wyatt E V., Diaz K, Griffin AJ, et al. Metabolic Reprogramming of Host Cells by Virulent Francisella tularensis for Optimal Replication and Modulation of Inflammation. J Immunol. 2016;196:4227–4236.

